# Comprehensive Complete-Genome Analysis of *Lactobacillaceae* and *Bifidobacteriaceae* Reveals Strain-Specific Metabolic Interactions in Chinese Gut Microbiota

**DOI:** 10.64898/2026.05.21.726744

**Authors:** Xin Tong, Hewei Liang, Yu Tian, Xinyu Yang, Ya Wang, Haoyu Wang, Yuzheng Gu, Zhihui Ma, Wenrui Su, Yanhong Liu, Sihang Cai, Zhiyin Lin, Peilin Zhang, Haifeng Zhang, Liang Xiao, Yiyi Zhong, Yuanqiang Zou

## Abstract

Bifidobacteriaceae and Lactobacillaceae are key probiotic families and widely used in food production, yet a comprehensive understanding of strain functions and their gut microbial interactions based on complete genomes remain understudied. Here we constructed a complete-genome dataset of 3,300 strains from these two families, including 1,151 newly isolated from China. Compared with draft assemblies, complete genomes substantially recovered a gene functional landscape encompassing stress tolerance, surface exopolysaccharide synthesis, nutrient utilization, and mobile genetic elements. Major species from both families exhibited a prevalence >60% in the Chinese population, far higher than that in US/Dutch cohorts. Notably, as a core probiotic species with remarkable genomic plasticity and gut-adaptive potential, *Lactiplantibacillus plantarum* stood out in our dataset for its enriched functional profile and was particularly abundant in the Chinese population. Moreover, compared with non-Chinese genomes, our isolates of key species displayed less metabolic complementarity and stronger competition with potentially pathogenic keystone species in the gut, thereby linking strain origin to enhanced probiotic potential and ecological fitness to benefit human gut health.

## Introduction

The gut microbiota plays a critical role in maintaining host health by regulating immune responses (Hou et al., 2022) and metabolic pathways (Choi et al., 2021). Among gut bacteria, members of Lactobacillaceae and Bifidobacteriaceae are widely recognized as beneficial to the host and are commonly used in probiotic products (Hutchinson et al., 2026). These bacteria can metabolize simple sugars such as fructose and lactose (Shao et al., 2026), as well as complex polysaccharides, including dietary fiber and human milk oligosaccharides (HMOs) (Shiver et al., 2025) (Lin et al., 2023). They can also inhibit pathogens by producing short-chain fatty acids (Fusco et al., 2023), bacteriocins, and other metabolites, while enhancing intestinal barrier function and modulating immune responses (Jaye et al., 2022).

Studies of probiotic functions have often relied on draft genomes. Large-scale genomic analyses have revealed the rich metabolic potential of lactic acid bacteria (LAB) (Jin et al., 2025), and accumulating high-resolution genomes have refined the taxonomy of *Bifidobacterium* (Shao et al., 2026). However, gaps and incomplete assemblies in draft genomes can cause key functional genes to be missed or falsely annotated, including biosynthetic gene clusters for secondary metabolites (Sanchez-Navarro et al., 2022) and multicopy genetic elements in complex genomic regions (Wang et al., 2026). Recent advances in long-read sequencing now make it possible to obtain complete genomes with high contiguity and completeness (Liang et al., 2025), enabling more comprehensive annotation of functional elements, including carbohydrate metabolism genes, antibiotic resistance genes, and related features (Xu et al., 2025) (Wang et al., 2026). Nevertheless, a systematic complete-genome database representing Lactobacillaceae and Bifidobacteriaceae from multiple sources and geographic backgrounds remains lacking.

Current microbiome research primarily focuses on species-composition changes induced by probiotic supplementation (Goswami et al., 2026) or on abundance differences across samples (Jin et al., 2025), whereas complementarity, competition, and functional metabolic networks between probiotics and other gut microbes remain poorly understood. Such interaction-level insights are essential for understanding how probiotics shape the gut ecosystem and, in turn, affect host health (Heinken et al., 2025). Notably, dietary habits, genetic backgrounds, and environmental exposures vary substantially across populations, potentially leading to divergent strain-specific interaction patterns in different geographic cohorts (Gu et al., 2026). Probiotics isolated from the gut, food, or environment may also exhibit distinct interaction modes with keystone gut microbes because of niche-specific adaptations (Deriu et al., 2013). These differences in interaction patterns may ultimately lead to distinct microbiota-modulating effects in healthy individuals and patients (Li et al., 2024).

Building on our previous culturomics work (Zou et al., 2019) (Lin et al., 2023), we targeted and cultured 784 Bifidobacteriaceae strains and 367 Lactobacillaceae strains from fecal samples of healthy Chinese individuals. We then performed long- and short-read sequencing followed by hybrid assembly to obtain complete genomes. We also retrieved 2,149 complete genomes from public databases after rigorous quality control, yielding a dataset of 3,300 complete genomes from Lactobacillaceae and Bifidobacteriaceae. Using this dataset, we constructed a comprehensive functional landscape of these strains and identified metabolic variation among isolates. Our analysis revealed distinct metabolic interaction patterns between strains from different geographic origins and keystone species in healthy and diseased individuals, providing a data foundation and analytical framework for precision probiotic interventions tailored to Chinese populations.

## Results

### Overall characteristics of the Bifidobacteriaceae and Lactobacillaceae genome dataset

We constructed a comprehensive dataset of 3,300 high-quality complete genomes from Bifidobacteriaceae and Lactobacillaceae, comprising strains isolated and sequenced in-house and strains downloaded from public databases. Figure 1 summarizes the global geographic distribution, phylogenetic relationships, and species-level composition of this dataset. In total, 1,151 strains were isolated in-house from human feces and yogurt samples collected in China. The remaining 2,149 public genomes were obtained from the NCBI and GTDB databases (downloaded March 10, 2026). All 3,300 genomes passed strict quality-control criteria: (i) genome completeness > 98% and contamination < 2%; (ii) N50 length greater than 90% of the total genome length; and (iii) fewer than 10 contigs.

**Figure 1.**
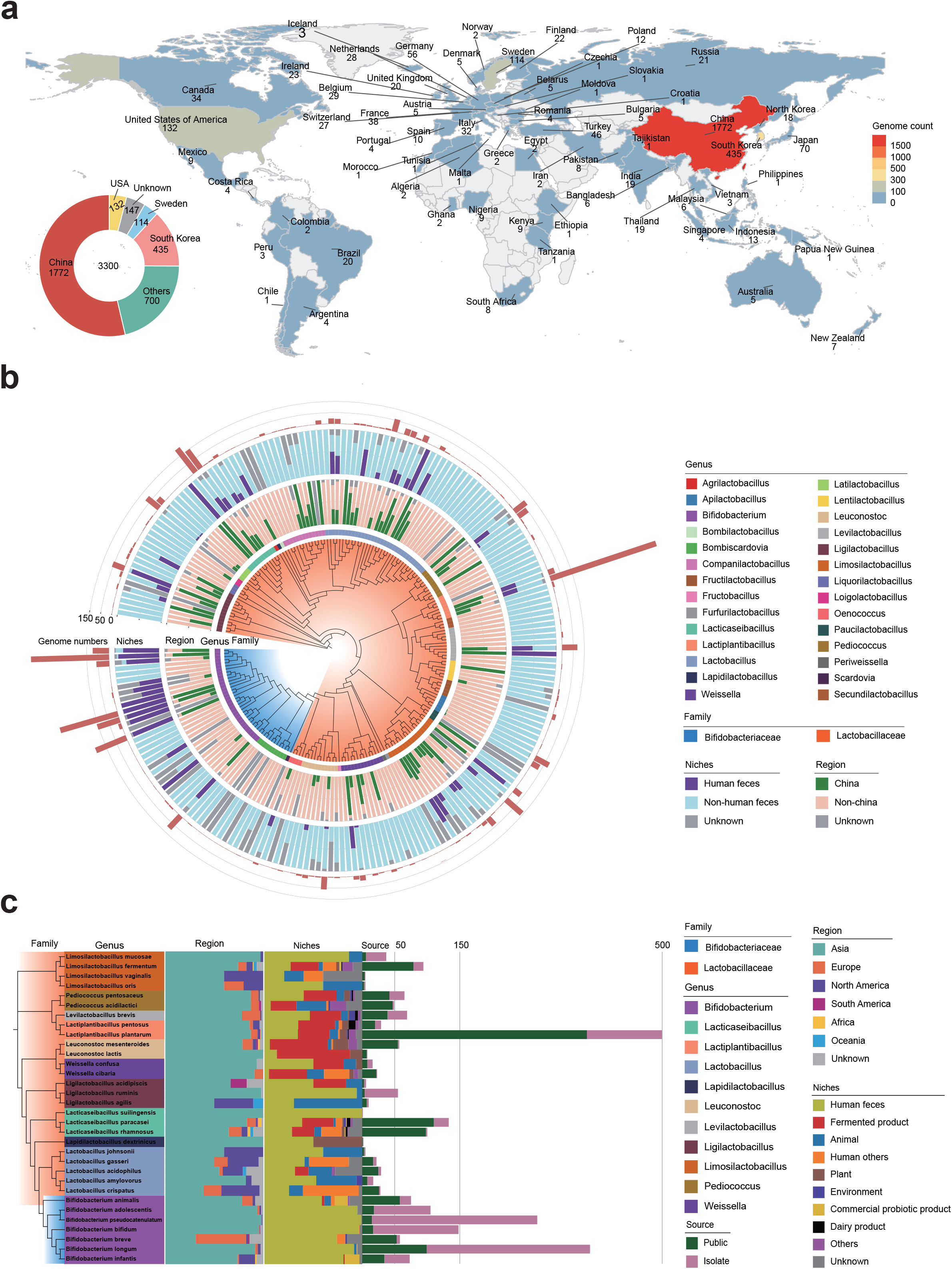

Figure 1a shows that the 3,300 genomes originated from a wide range of geographic locations across six continents, with Asia accounting for the largest proportion. The 1,151 in-house isolates represented a substantial fraction of the Asian genomes and markedly increased the representation of Chinese strains (Figure 1b). China contributed the largest number of genomes worldwide, with the in-house isolates forming the core component, followed by North America (United States, 152 genomes), Europe (Sweden, 114 genomes), and South America (Brazil, 20 genomes). This distribution indicates that the in-house collection helped fill gaps in regional genomic resources and reduced geographic imbalance in strain representation. Chinese-origin strains were densely distributed across branches of multiple key genera, highlighting the contribution of our isolates to East Asian taxonomic coverage. Human-feces-derived strains were concentrated in specific clades, including *Bifidobacterium* and *Lactobacillus*, whereas strains from non-human-fecal sources were broadly distributed across Lactobacillaceae. This pattern provides a basis for subsequent comparisons between gut and non-gut strains and between Chinese and non-Chinese strains. At the species level, the in-house strains substantially increased genomic diversity in key taxa with probiotic or industrial relevance, including *Lactiplantibacillus plantarum, Bifidobacterium longum*, and *Bifidobacterium pseudocatenulatum* (Figure 1c). This contribution was further supported by the pan-genome accumulation curves (Figure S1): for most species, curves incorporating the in-house strains consistently lay above those based only on public genomes, indicating that our isolates harbor novel gene content and expand the known pan-genome.

### Complete genomes recover assembly-sensitive probiotic functions missed in draft assemblies

To determine whether genome completeness affects strain-level functional inference, we compared functional annotations between complete genomes and available draft assemblies in the probiotic genome resource. The complete-genome collection contained 3,300 unique strains, whereas the draft-genome collection contained 1,149 unique strains, with 793 strains represented in both resources. After filtering to high-confidence paired strain-gene comparisons, 417 strains and 334,952 strain-gene combinations were retained for direct count-based analysis.

The transition from draft to complete assemblies significantly improved genome continuity and the recovery of key genomic features. As shown in Figure S1A, complete genomes had significantly higher N50 values and recovered many ribosomal RNA (rRNA) genes and CRISPR arrays that were truncated or missing in draft assemblies. Although 99.75% of gene annotations remained unchanged after excluding uncharacterized categories and outliers, complete genomes retained a clear excess of positive differences (713 entries) over negative differences (123 entries), resulting in a net gain of 1,099 annotated hits. At the strain level, 412 of the 417 paired strains showed a positive aggregate difference.

Genes preferentially recovered in complete genomes were concentrated in repeat-rich genomic compartments that are typically difficult to resolve in fragmented assemblies. Complete genomes showed a marked increase in the detection of insertion sequences (IS), with the IS30, IS3, and ISL3 families showing the largest gains (Figure S1b-d). Analysis across 23 species and 793 genomes revealed that these gains primarily involved mobile genetic elements, defense/resistance systems, and metabolic regulation (Figure 2a). Notable recovery was observed in core probiotic genera such as *Bifidobacterium, Lactobacillus*, and *Lacticaseibacillus*, particularly for genes such as istB, hsdM, and diverse transposases (Figure 2a).

**Figure 2.**
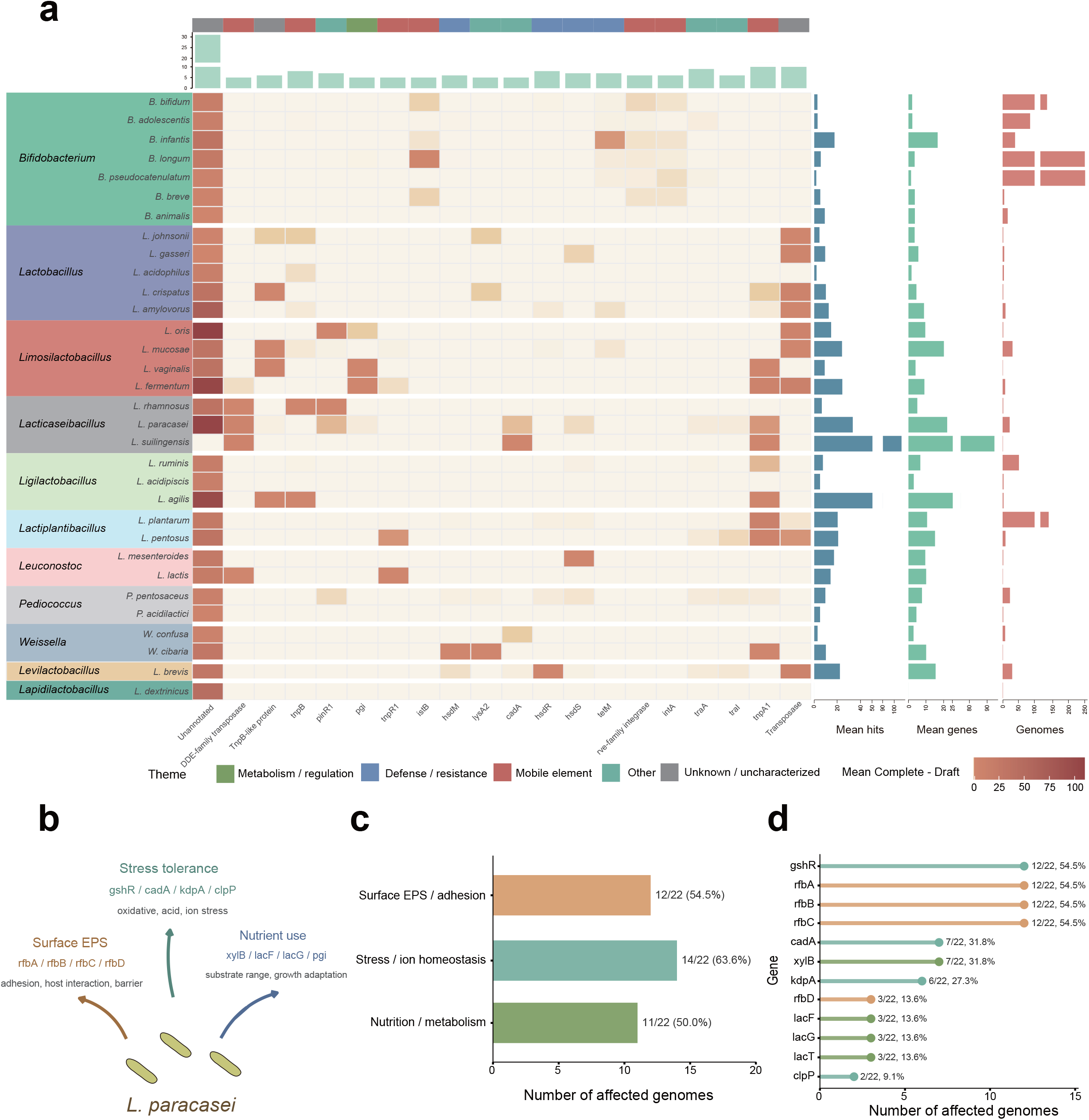

Several recurrent genes illustrated how complete genomes changed biologically interpretable probiotic annotations, particularly in *Lacticaseibacillus paracasei* (Figure 2b). Complete assemblies recovered essential genes associated with stress tolerance (gshR, cadA, kdpA, clpP), surface exopolysaccharides (EPS; rfbA/B/C/D), and nutrient utilization (xylB, lacF/G/T) (Figure 2b). Quantitatively, 63.6% of affected genomes showed corrected annotations for stress and ion homeostasis, while more than 50% showed improvements in surface EPS/adhesion and nutrition/metabolism functions (Figure 2c, d). Genes such as gshR and the rfb cluster were recovered in more than 50% of the relevant strains (Figure 2d). These findings underscore the need to use complete genomes as reference-standard resources for downstream analyses of probiotic function, safety, and ecological compatibility.

### Source-associated differences in accessory genomes and putative probiotic functions

Using Roary pan-genome presence/absence matrices, we compared accessory gene composition between human gut-derived and non-human-gut-derived strains within each species. The species-level functional enrichment heatmap showed that several species contained gene clusters enriched in human gut-derived strains, mainly involving carbohydrate and host-glycan utilization, cell-surface structures, EPS and cell-envelope remodeling, stress tolerance, transport, and regulatory functions (Figure 3a). PCoA based on contrast-informative accessory genes further revealed visible separation between human gut-derived and non-human-gut-derived strains in selected species (Figure 3b), suggesting that strain source was associated with accessory genome composition.

**Figure 3.**
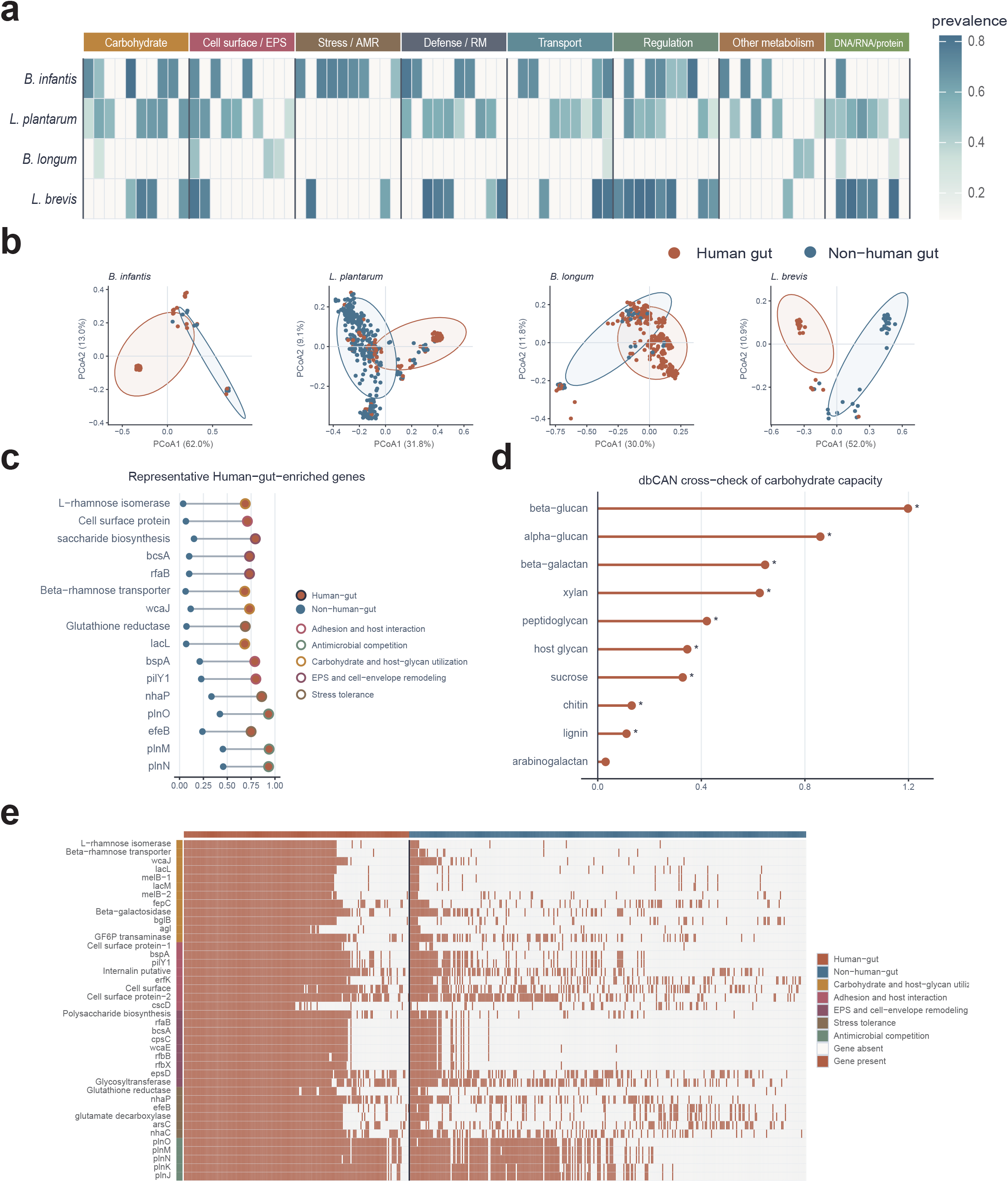

In the representative species *L. plantarum*, human gut-derived strains were enriched for genes potentially related to gut adaptation and probiotic-associated functions, including genes involved in carbohydrate transport and metabolism, cell-surface proteins, polysaccharide biosynthesis, stress-associated functions, and plantaricin-related modules (Figure 3c). The strain-level heatmap showed that these genes occurred at higher prevalence across a substantial proportion of human gut-derived strains, rather than being driven by only a few individual genomes (Figure 3E). In addition, dbCAN-based cross-validation showed that human gut-derived *L. plantarum* strains had greater genomic coding potential for carbohydrate-active enzymes associated with substrates such as beta-glucan, alpha-glucan, beta-galactan, and xylan (Figure 3d). Together, these results indicate that accessory genomes of human gut-derived strains are enriched for functional modules potentially linked to adaptation to the gut environment.

Because human gut-derived strains were collected from different geographic backgrounds, we further compared Chinese and non-Chinese human gut-derived strains. This stratified analysis showed a weaker signal than the gut-versus-non-gut comparison, but accessory genome separation remained detectable in selected species (Figure S2a). In *L*.*plantarum*, genes enriched in Chinese gut-derived strains were mainly associated with carbohydrate and host-glycan utilization, adhesion and host interaction, and EPS and cell-envelope remodeling modules (Figure S2b). dbCAN annotations further supported differences in the predicted carbohydrate-active enzyme potential of Chinese gut-derived *L. plantarum* strains (Figure S2c). Thus, geographic background may contribute additional accessory genome variation within human gut-derived strains, although this signal was more limited than the primary source-associated comparison.

Acid tolerance is crucial for probiotic functionality. We therefore cultured 120 Bifidobacteriaceae and 73 Lactobacillaceae strains for 24 h at pH 3.0 and pH 2.0. After 24 h, OD values increased for most strains, with only 13 and 22 strains showing decreased OD values at pH 3.0 and pH 2.0, respectively. Substantial variation in OD changes was observed among strains (Figure S3a). To identify key genes underlying differential acid tolerance, we applied a logistic regression model to extract relevant features using 5 x 5 cross-validation. For Bifidobacteriaceae, the mean AUC was 0.700 +/- 0.091 at pH 2 and 0.812 +/- 0.102 at pH 3. For Lactobacillaceae, the mean AUC was 0.805 +/- 0.117 at pH 2 and 0.747 +/- 0.138 at pH 3 (Figure S3b). Among the top 10 genes from these models (Figure S3c), four were related to ABC transporters and five to membrane transport, suggesting that cation efflux is a key factor for acid tolerance in both Bifidobacteriaceae and Lactobacillaceae.

### Geographic prevalence and strain--specific metabolic advantages of Lactobacillaceae and Bifidobacteriaceae

Consistent with geographic strain specificity, the abundance and prevalence of gut microbial species have also been reported to vary by region. Accordingly, we analyzed data from the Chinese Gut Microbial Reference (CGMR), the Human Microbiome Project (HMP) cohort, and a Dutch metagenomic cohort. Gut microbial communities showed significant biogeographic differentiation among Chinese, American, and Dutch populations, with similar divergence in the composition of Lactobacillaceae and Bifidobacteriaceae species (Figure 4a, b). PERMANOVA (R2 = 0.129 and 0.126, both p = 0.001) and PCoA confirmed robust country-level effects and distinct clustering by nationality, indicating host-geography-dependent signatures.

**Figure 4.**
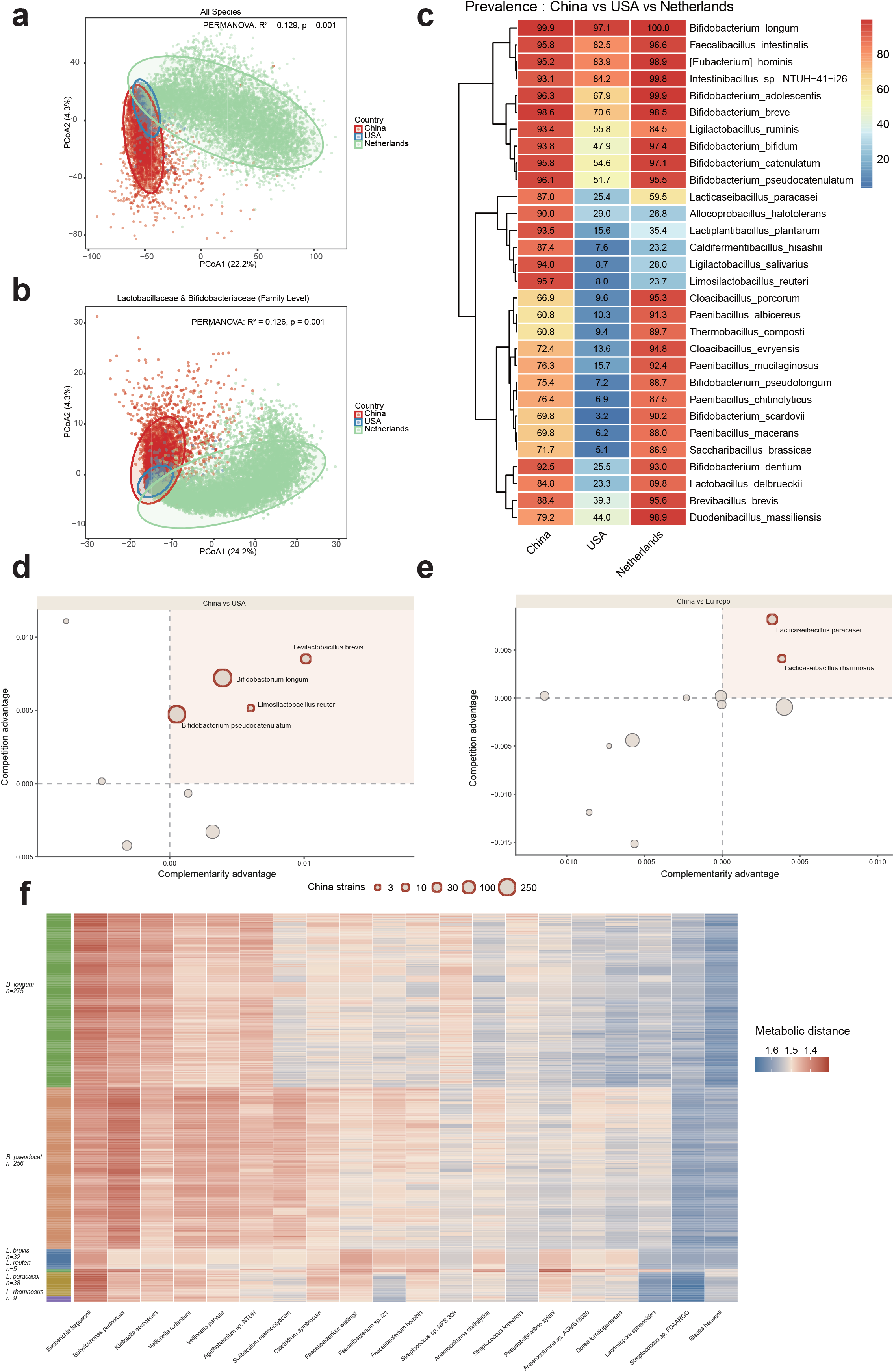

Based on the top 30 prevalent species of Lactobacillaceae and Bifidobacteriaceae, these taxa could be grouped into three clades (Figure 4c). The first clade, consisting of 10 species including *Bifidobacterium longum* and *Faecalibacillus intestinalis*, showed relatively high prevalence across all three countries. The second clade, comprising six species such as *Lacticaseibacillus paracasei* and *Lactiplantibacillus plantarum*, was predominantly detected in the Chinese cohort but rarely observed in the American and Dutch cohorts. The remaining 14 species, represented by *Bifidobacterium pseudolongum* and *Bifidobacterium dentium*, showed high prevalence in both the Chinese and Dutch cohorts but low prevalence in the American cohort. Overall, the prevalence of Lactobacillaceae and Bifidobacteriaceae species in China exceeded 60%, a level considerably higher than that in the American (HMP) cohort, suggesting that members of these families are more common in the Chinese population and may play a more prominent role in the gut microbiota.

To further investigate the potential influence of Lactobacillaceae and Bifidobacteriaceae on the gut microbiota, we constructed a co-occurrence network based on the CGMR cohort. From network connectivity, we identified the top 20 keystone species and performed metabolic interaction analysis with Lactobacillaceae and Bifidobacteriaceae species for which complete genomes were available. Among these species, *Levilactobacillus brevis, Bifidobacterium longum, Limosilactobacillus reuteri*, and *Bifidobacterium pseudocatenulatum* showed stronger complementarity and competition advantages with keystone species in the Chinese cohort than did genomes from the United States (Figure 4d, e). Notably, *Bifidobacterium longum* and *Bifidobacterium pseudocatenulatum* were also the species from which we isolated the largest number of strains. Compared with European genomes, *Lacticaseibacilus paracasei* and *Lacticaseibacilus rhamnosus* also showed distinct metabolic interaction advantages. Based on the metabolic models, these species had short metabolic distances to *Escherichia fergusonii, Butyricimonas paravirosa*, and *Klebsiella aerogenes* (Figure 4f), suggesting potentially frequent metabolic interactions with these species. Among them, *Bifidobacterium pseudocatenulatum* showed short metabolic distances to most of the top 20 keystone species, except *Streptococcus* sp. FDAARGO and *Blautia hansenii*.

### Metabolic interactions of Lactobacillaceae and Bifidobacteriaceae with disease--associated keystone species across multiple patient cohorts

Many probiotics, particularly those belonging to Lactobacillaceae and Bifidobacteriaceae, benefit patients with various diseases (Quigley and Shanahan, 2025). To investigate the metabolic interaction patterns of these probiotic taxa, we analyzed cohorts of patients with colorectal cancer (CRC) (Gao et al., 2022), obesity (OB) (Liu et al., 2017), hyperlipidemia (HTN) (Yan et al., 2017), non-alcoholic fatty liver disease (NAFLD) (Leung et al., 2022), and inflammatory bowel disease (IBD; two cohorts) (Lloyd-Price et al., 2019). Overall, the composition of Bifidobacteriaceae and Lactobacillaceae did not differ markedly between disease and healthy control samples (Figure 5a). Based on genus-level relative abundances, *Limosilacyobacillus, Fructilactobacillus* and *Bifidobacterium* exhibited higher z-scores across multiple control groups, indicating that these genera were generally more abundant in healthy individuals (Figure 5b).

**Figure 5.**
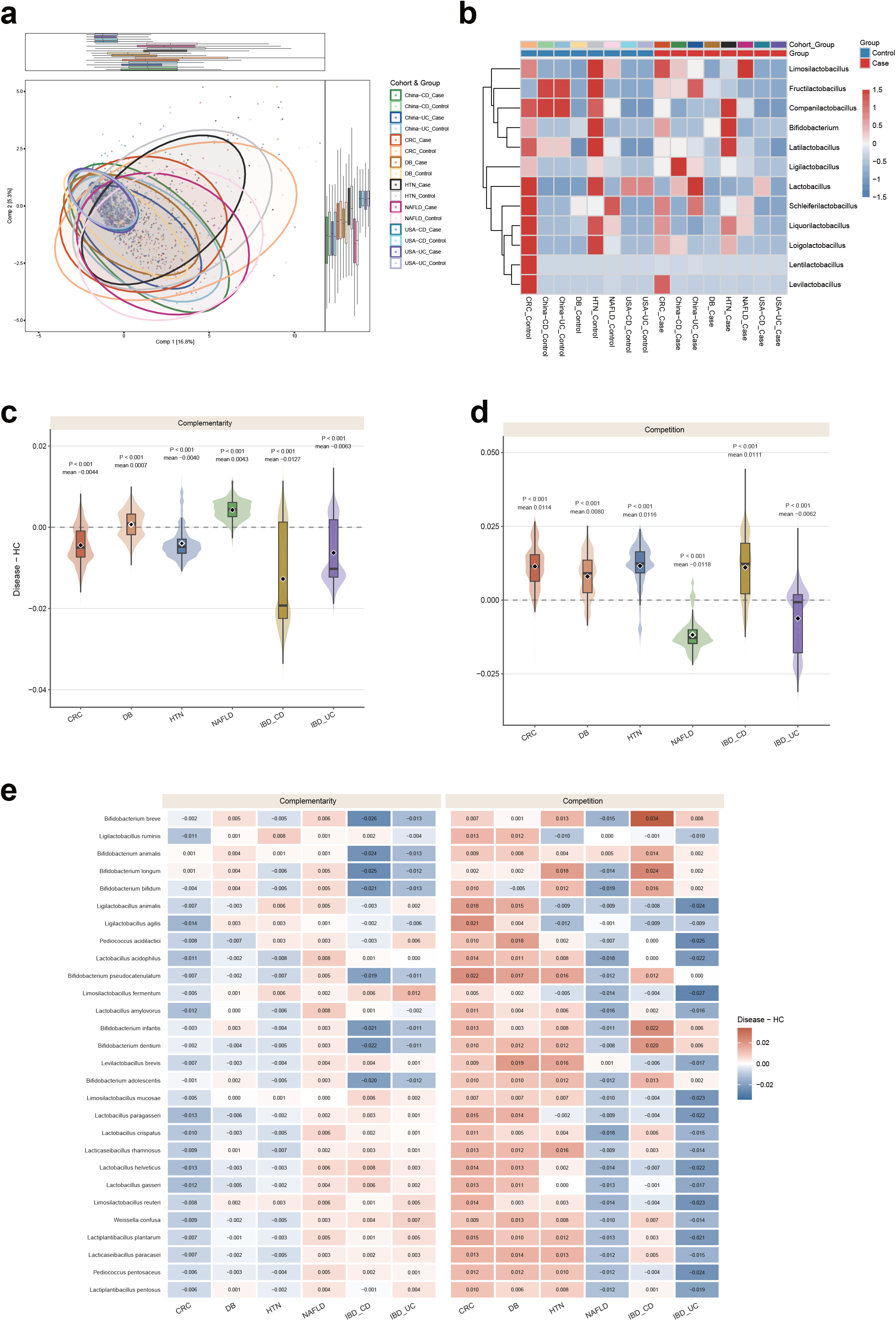

Overall, Lactobacillaceae and Bifidobacteriaceae species showed weaker complementarity and stronger competition with disease-associated keystone species in the CRC, HTN, and CD cohorts (Figure 5c, d), suggesting that they may compete with keystone species in the gut microbiota of these patients and potentially inhibit their growth. Most Lactobacillaceae and Bifidobacteriaceae species showed consistent patterns across these three cohorts (Figure 5e). Notably, *Bifidobacterium breve* showed higher competition and lower complementarity values in the CD cohort than other species, indicating that this species may play a more substantial role in modulating the gut microbiota at the metabolic-interaction level.

To investigate whether the origin of Lactobacillaceae and Bifidobacteriaceae strains influences their metabolic interactions with the gut microbiota, we compared gut-derived and non-gut-derived strains. We identified 12 species in which gut-derived strains exhibited superior complementarity and competition advantages with keystone species from healthy individuals compared with non-gut-derived strains. Among these, *Levilactobacillus brevis, Bifidobacterium longum* and *Lacticaseibacilus paracasei*--species from which we isolated the largest numbers of strains--all showed short metabolic distances to most gut keystone species.

## Discussion

The construction of a comprehensive dataset comprising 3,300 high-quality genomes substantially expands the genomic resources available for Bifidobacteriaceae and Lactobacillaceae. Although public databases contain valuable resources for these families, they are dominated by strains from Western populations and industrial settings, with limited representation of isolates from East Asian, particularly Chinese, human gut and fermented food sources. By integrating 1,151 newly isolated and sequenced strains from Chinese human feces and yogurt samples with 2,149 publicly available genomes, this study broadens the taxonomic, geographic, and ecological coverage of available genomic data. In this dataset, China became the largest contributor of genomic resources, with our in-house strains forming the core component. This collection substantially increases both the number and diversity of Chinese-origin strains and enriches the representation of gut-associated and fermented-food-derived isolates, which are of particular interest for probiotic and functional food research. The phylogenetic distribution further underscores this contribution: Chinese-origin strains are densely distributed across key genera, and human fecal isolates form distinct concentrated clades, providing a robust framework for comparisons between gut and non-gut strains and between Chinese and non-Chinese strains.

At the species level, the impact of our in-house strains was particularly pronounced for taxa of high probiotic and industrial relevance, such as *Lactiplantibacillus plantarum, Bifidobacterium longum*, and *Bifidobacterium pseudocatenulatum*. Pan-genome analyses indicate that differences between human gut-derived and non-human-gut-derived strains were mainly reflected in the accessory genome. Compared with core genes, accessory genes are more likely to capture adaptation to specific ecological niches. The enrichment of genes related to carbohydrate utilization, cell-surface interaction, EPS and cell-envelope remodeling, and stress tolerance in human gut-derived strains suggests that these strains may have greater potential for adaptation to the gut environment. The focused analysis of *L. plantarum* further supported this interpretation, particularly through enrichment of carbohydrate-utilization and cell-surface-associated genes and dbCAN-supported differences in carbohydrate-active enzyme coding potential.

Within human gut-derived strains, Chinese and non-Chinese isolates also showed some accessory genome stratification, although this difference was weaker than that between gut and non-gut sources. This result should therefore be interpreted as supplementary evidence that geographic background may contribute to strain-level functional variation, rather than as a deterministic geographic effect. Because this analysis is based on genome-derived functional predictions, it reflects functional potential rather than direct phenotypic activity. The presence of a gene does not necessarily indicate that it is expressed or functional in the gut environment, and differences in public database composition, sampling region, and isolation background may introduce bias. Further transcriptomic, metabolomic, and phenotypic validation will be required to determine whether these candidate functions contribute to measurable probiotic traits.

Our results reveal distinct biogeographic patterns in Lactobacillaceae and Bifidobacteriaceae distribution, with Chinese populations showing higher prevalence (>60%) of these taxa compared to Western cohorts. Notably, species like *Lacticaseibacillus paracasei* and *Lactiplantibacillus plantarum* demonstrated China-specific enrichment, while *Bifidobacterium pseudolongum* showed shared prevalence in Chinese and Dutch populations. These geographic patterns were accompanied by functional differences - Chinese isolates of *Bifidobacterium longum* and *Bifidobacterium pseudocatenulatum* exhibited stronger metabolic complementarity with local keystone species compared to Western strains.

Lactobacillaceae and Bifidobacteriaceae species actively compete with disease-associated keystone species in CRC, HTN and CD patients, suggesting their potential to suppress pathogenic microbiota development through niche competition. The particularly strong competitive effect of *Bifidobacterium breve* in CD patients highlights its promising therapeutic value for IBD treatment. These consistent competitive patterns across multiple disease cohorts reveal an important ecological mechanism: probiotic strains may exert beneficial effects not only through direct host modulation, but also by outcompeting pathogenic species for ecological niches and resources. This competitive exclusion mechanism could represent a key strategy for rebalancing gut microbiota in various disease states. Future studies should focus on identifying the specific competitive factors (e.g., nutrient utilization advantages, antimicrobial production) that enable these probiotic species to effectively suppress disease-associated microbes.

## Materials and Methods

### Complete genome dataset of Bifidobacteriaceae and Lactobacillaceae

The sample collection, strain isolation and culture, and analysis were approved by the Institutional Review Board on Bioethics and Biosafety of BGI under th e numbers BGI-IRB 22112 and BGI-IRB 26021.

The complete genome dataset of Bifidobacteriaceae and Lactobacillaceae constructed in this study comprises publicly available genomes and genomes of strains independently isolated by our research group. We downloaded all complete genomes of Bifidobacteriaceae and Lactobacillaceae submitted between January 1, 2015, and March 10, 2026, from the NCBI and GTDB databases (https://www.ncbi.nlm.nih.gov/; https://gtdb.ecogenomic.org/). Strains sequenced in-house we re isolated and purified by our group, and whole-genome sequencing was outsourced to Shenzhen BGI. Sequencing was performed on the MGI DNBSEQ-T1 short-read platform in paired-end mode (PE150) and complemented by the MGI CycloneSEQ long-read platform to obtain long fragments. Raw sequencing dat a were uniformly subjected to quality control and preprocessing: short reads we re quality-filtered and adapter-trimmed using fastp (v0.20.0) (Chen, 2025), and long reads were filtered for low-quality sequences using NanoFilt (chopper) (D e Coster et al., 2018). The preprocessed short- and long-read data were assembled using Unicycler (v0.5.1) in a hybrid approach to generate high-quality com plete genomes (Wick et al., 2017). All genomes were quality-controlled using CheckM2 (v1.1.0) with the uniref100.KO.1.dmnd database according to the foll owing criteria: (1) genome completeness > 98% and contamination < 2%; (2) N50 length > 90% of the total genome length; and (3) fewer than 10 contigs (Chklovski et al., 2023). In addition, all genomes were taxonomically annotated using GTDB-Tk (v2.5.2) with the GTDB R226 database, and only genomes strictly identified as Bifidobacteriaceae or Lactobacillaceae were retained (Chaum eil et al., 2022).

### Draft genome assembly

For strains independently isolated by our research group, draft genome assembly was performed using raw paired-end short-read sequencing data. Raw reads were first subjected to quality control using fastp (v0.20.0), including quality filtering, adapter trimming, and removal of reads shorter than 30 bp (Chen, 2025). The resulting high-quality reads were then assembled using Unicycler (v0.5.1) with default parameters, yielding a draft genome sequence for each strain (Wick et al., 2017).

### Phylogenetic tree construction

Representative genomes were selected for each species by maximizing the value of completeness - 5 x contamination across all genomes of that species (Olm et al., 2017). Phylogenetic trees were then constructed for all representative genomes using GTDB-Tk (v2.5.2) with the GTDB R226 database. First, multiple sequence alignment was performed using the GTDB-Tk align module, generating an alignment file in .msa.fasta format. Based on this alignment file, the gtdbtk infer command was used to build a species tree with the maximum likelihood method (Chaumeil et al., 2022).

### Pan-core genome construction

Based on the GFF3-format annotation files generated by Prokka, the pan-genome and core genome for each species were constructed using Roary (Seemann, 2014) (Page et al., 2015). Pan-genome analysis was performed for species represented by more than three genomes in both public databases and our independently sequenced dataset. Core genes were defined as genes present in 99%-100% of strains, whereas the pan-genome was defined as all genes present in 0%-100% of strains.

### Pan-genome-based differential gene enrichment and annotation

Genome functional annotation was performed using Bakta v1.12.0 (Schwengers et al., 2021) with default parameters, and carbohydrate-active enzymes were annotated using dbCAN v4.1.4 (Zheng et al., 2023) in protein mode. Based on the previously generated Roary gene presence/absence matrices, we compared accessory gene composition between strains from different sources within each species. The primary comparison was performed between human gut-derived and non-human-gut-derived strains. A secondary stratified analysis was restricted to human gut-derived strains and compared Chinese and non-Chinese isolates. Strain source metadata were integrated with the gene presence/absence matrices, and only species with at least three strains in both comparison groups were retained. Gene prevalence was calculated separately within each group. Differential gene enrichment was assessed using Fisher’s exact test followed by Benjamini-Hochberg false discovery rate correction. Genes were considered significantly enriched when FDR < 0.05, absolute prevalence difference >= 0.20, enriched-group count >= 3, and enriched-group prevalence >= 0.05. Significantly enriched genes were used to generate species-level functional enrichment heatmaps and representative gene plots, whereas contrast-informative accessory genes were used for PCoA based on Jaccard distance. Enriched genes in the representative species *L. plantarum* were further interpreted using post hoc functional annotation and dbCAN-based carbohydrate-active enzyme profiles.

### Construction of a prediction model for acid--base tolerance

Using a gene presence/absence matrix from Roary (Page et al., 2015) pan-genome analysis of Bifidobacteriaceae and Lactobacillaceae strains, we performed binary classification for high tolerance (growth > 0.2) versus low tolerance (growth <= 0.2). Features were filtered by frequency (15%-85%), normalized, and selected via recursive feature elimination with cross-validation (RFECV). A logistic regression model (L2 regularization, balanced class weights) was evaluated using 5 x 5 repeated stratified cross-validation, and gene importance was derived from regression coefficients.

### Estimation of species abundance and prevalence

Taxonomic classification was performed using Kraken 2 (v2.16.0) (Wood et al., 2019) with the k2_pluspfp_20251015 database, followed by abundance estimation with Bracken (v3.0.1) (Lu et al., 2017). For each sample, a taxon was considered present if the estimated read count assigned to that taxon exceeded 100. The relative abundance of each bacterial species was calculated as the estimated read count assigned to that species divided by the total estimated read count assigned to all bacterial species. These procedures were applied uniformly to all cohorts in this study.

### GEM reconstruction and metabolic interaction analysis

Genome-scale metabolic models (GEMs) for all gut-derived genomes were reconstructed using CarveMe (v1.5.1) with customized parameters (Machado et al., 2018). Unlike conventional bottom-up approaches that require manual curation and predefined growth media, our workflow incorporated gut-relevant culture media and gap-filling to ensure model completeness. The input for CarveMe consisted of coding sequences predicted with Prokka, and pairwise interactions between GEMs were subsequently analyzed using PhyloMint (Lam et al., 2020).

### Network construction and identification of keystone nodes

FastSpar (Watts et al., 2019) was used to perform SparCC co-occurrence network construction for each sample, generating edge lists with correlation coefficients. Nodes represented microbial taxa, and edges were undirected, with absolute correlation used as the weight. For each network, node topological influence was quantified using the Influential (IVI) index (R package influential, d = 3) (Salavaty et al., 2020). To identify keystone nodes, nodes were removed in descending IVI order, and the resulting decline in largest connected component size was compared against 1,000 random node-removal permutations. Nodes with p < 0.001 were considered keystone nodes. For each network, the top 20 keystone nodes by IVI value were retained for downstream analysis. All computations were performed in R (version 4.2.2) using packages influential, igraph, and dplyr.

## Acknowledgements

This work was supported by the Shenzhen Municipal Government of China (No. JCYJ20241202124801003 and KCXFZ20240903094006009) and the Shenzhen Science and Technology Program (No. SYSPG20241211173845014). We thank our colleagues at BGI Research for their assistance with strain cultivation, DNA extraction, library construction, and sequencing. We also thank DCS Cloud (https://cloud.stomics.tech/) for providing the computational resources and software support necessary for this study.

## Contributions

Conceived and designed the experiments: Y. Zou, Y. Zhong, L. X., and X. T. Performed the experiments: Z. M., W. S., Y. L., S. C., Z. L., and P. Z. Analyzed the data: X. T., H. L., Y. T., X. Y., Y. W., H. W., and Y. G. Contributed reagents/materials/analysis tools: H. Z., L. X., and Y. Zhong. Wrote the paper: H. L., Y. T., X. Y., Y. W., H. W., and Y. Z. All authors commented on the manuscript.

